# Physical Exercise Improves Working Memory through Ripple-Spindle Coupling

**DOI:** 10.1101/2024.07.10.602896

**Authors:** Xinyun Che, Benedikt Auer, Paul Schmid, Christoph Reichert, Annemarie Scholz, Tom Weischner, Robert T. Knight, Stefan Dürschmid

## Abstract

Spindle-ripple coupling enhances memory consolidation during sleep. Ripples, representing the compressed reactivation of environmental information, provide a mechanism for retaining memory information in chronological order and are also crucial for working memory (WM) during wakefulness. Brief sessions of physical exercise (PE) are proposed to boost WM. In concurrent EEG/MEG sessions, we investigated the role of PE in WM performance and high-frequency-ripple to spindle coupling. Ripples, identified in MEG sensors covering the medial temporal lobe (MTL) region, predicted individual WM performance. Ripples were locked to robust oscillatory patterns in the EEG defined spindle band. Spindle activity and ripples decrease during initial stimulus presentation and rebound after 1 sec. Behaviorally, PE enhanced WM performance. Neurophysiologically, PE scaled the ripple rate with the number of items to be kept in WM and strengthened the coupling between ripple events and spindle oscillations. These findings reveal that PE enhances WM by coordinating ripple-spindle interaction.

## Introduction

Working memory (WM) allows information to be stored in a sequential order for future retrieval. A common method to test working memory capacity in humans is the N-back task requiring subjects to match items that appeared in previous trials. In the N-back task, target detection performance decreases with an increasing amount of information to be retained due to the finite capacity of WM. Brief sessions of physical exercise (PE) enhance working memory performance^1–3^. However, the neurophysiological mechanism for retaining information in chronological order within WM and the impact of PE on this mechanism remain unclear. Sleep research suggests a specific interaction between high-frequency ripples and thalamic generated spindle activity as a key mechanism for successful integration of information into memory^4–6^. High-frequency ripple events, manifesting as transient bursts (~80–150 Hz in humans^7^), are considered a compressed reactivation of sequential environmental information during cued recall. Spindle activity is assumed to establish a time frame for ripples occurrence^7^. However, whether ripple-spindle coupling during wakefulness is a mechanism for memory formation is unknown.

In simultaneous EEG/MEG recordings, we assessed whether performing an N-back WM task is tracked by modulation in spindle activity, ripple rate and spindle-ripple coupling. Specifically, we tested whether PE improved both WM capacity and MEG-ripple – EEG-spindle coupling. In rest session, participants remained inactive in short experimental breaks (2 min) of typical EEG/MEG recordings. In PE session, participants were engaged in movement during breaks (2 min) using an MEG-compatible pedal trainer designed to facilitate independent forward and backward movements, resembling a walking motion.

PE enhanced target detection. High-frequency ripples, key for organizing information into working memory, were detected in MEG sensors covering the medial temporal lobe (MTL) region. Both EEG-spindle and MEG-ripple rates decreased during stimulus presentation but increased after 1 sec. EEG-spindle activity, a key marker for memory consolidation in sleep research, was higher in the PE session compared to rest during rebound. The rebound MEG-ripple rate predicted individual WM performance and scaled with the number of items retained in WM. Critically, PE increased coupling of MEG-ripples to EEG-spindle activity also enhanced WM performance.

## Results

### Procedure

21 participants participated in the experiment with simultaneous EEG and MEG recording. The experiment was conducted in two sessions, with and without PE, on two different days. The order of PE and rest was counterbalanced across subjects. Each session consisted of 12 blocks. During breaks between blocks subjects were instructed to rest or use an MEG compatible pedal trainer for two minutes. The pedal trainer permitted independent forward and backward movements (see ***Fig. 1A*** and ***Methods***). Participants were directed to pedal at a moderate speed, adjusting their pace individually, akin to a walking motion. Within each block, an item (one of the 26 letters of the Latin alphabet) was a target if it matched the item from N=2, N=3, or N=4 trials ago (N-back task), requiring subjects to hold a varying number of items in working memory in each block (see ***Fig. 1A*** and ***Methods*** for a detailed description of the procedure).

**Fig. 1.**
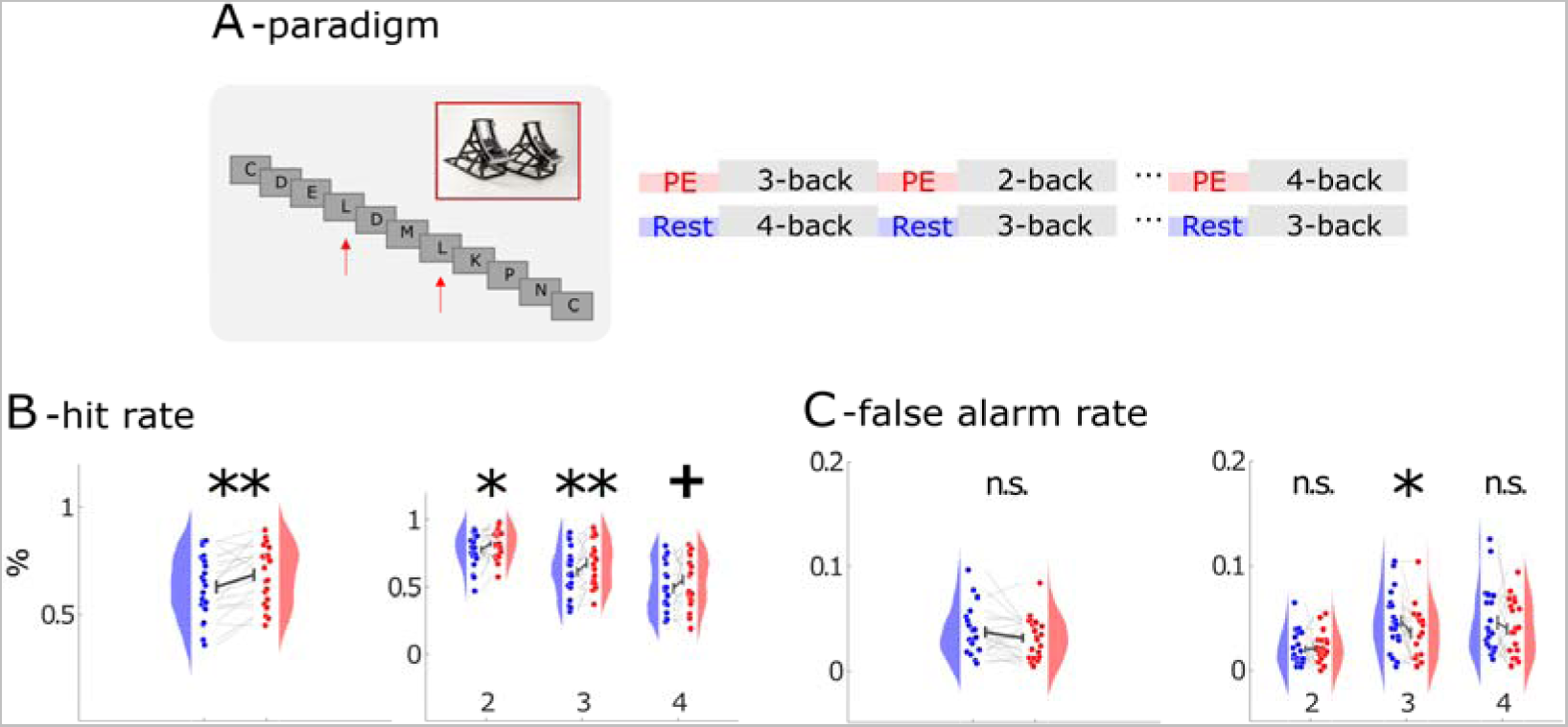
Behavioral Performance. ***A***: Task design. Subjects were instructed to rest or use a pedal trainer before the N-back task. ***B***: (left): shows that hit rate increased across all conditions. (right): Hit rate across different N-back conditions. ***C***: (left): Individual false alarm rate. (right): False alarms across different N-back conditions. Red indicates PE and blue rest session.

#### I. Behavioral performance

First, we evaluated target detection accuracy (hit rate; H; see ***Fig. 1B***) and false alarms (see ***Fig. 1C***). When we compared hit rate, we found a main effect of PE (F_(1,120)_ = 4.04; p = 0.047) and N-back condition (F_(2,120)_ = 31.7; p < .0001). Performance declined with number of items to be held in working memory (H_2_ = 80.07%; H_3_ = 64.68%; H_4_ = 52.12%). However, hit rate was higher after PE than after rest (H_PE_ = 68.51%; H_rest_ = 62.74%; t_20_ = 3.62; p = .0017) in all N-back conditions (2-back: t_20_ = 2.78; p = 0.012; 3-back: t_20_ = 3.39; p = 0.003; 4-back: t_20_ = 2.21; p = 0.039, see ***Fig. 1B***). When we compared false alarm rates, except for the 3-back condition (t_20_ = 2.55; p = 0.019), no significant effect of PE was found(F_(1,120)_ =1.56; p=0.214, see ***Fig. 1C***).

#### II EEG spindle activity

Spindle activity provides a time frame for ripple occurrence^7^. In the EEG, thalamic spindle activity is maximal at fronto-central electrodes^8^. We found a consistent temporal evolution across both rest and PE with the spindle activity decreasing immediately after stimulus onset, followed by a subsequent increase in fronto-central EEG electrodes (see ***Fig. 2A*** left) peaking at 1150 msec after stimulus presentation with enhanced activity between 1 to 2 sec after stimulus presentation following PE (t_20_ = 2.20; p = 0.04).

**Fig. 2.**
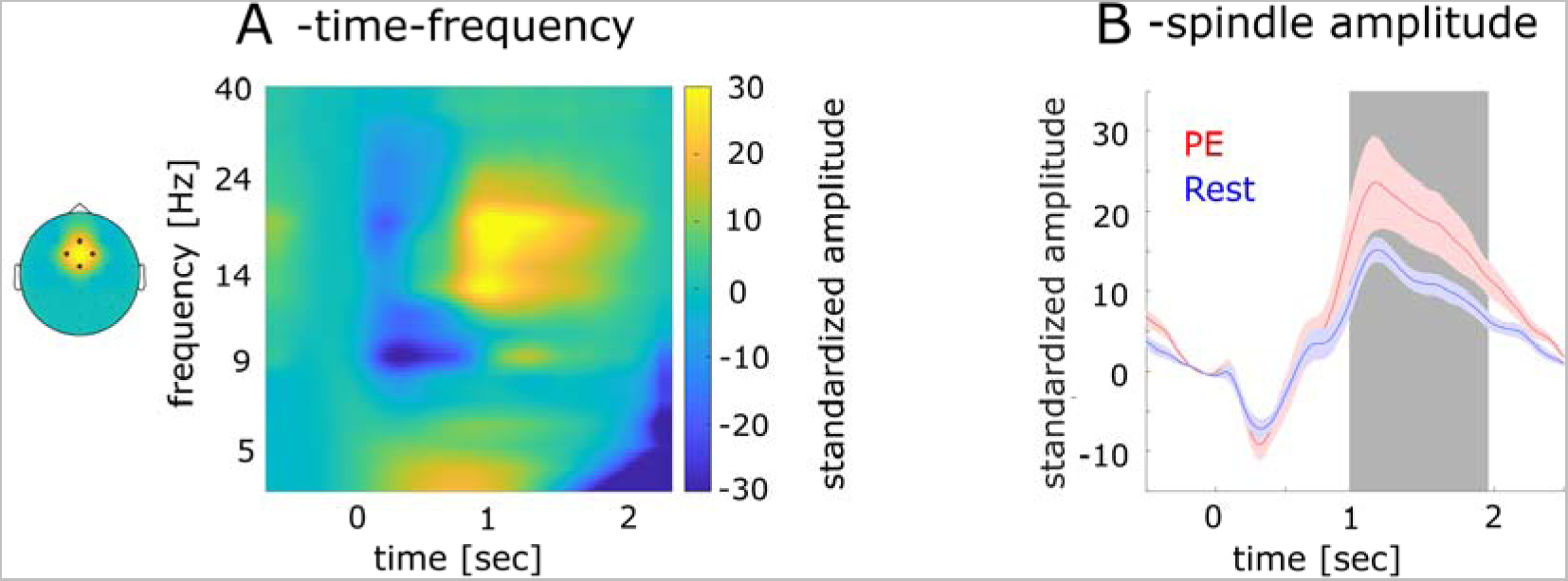
Frequency Amplitude of EEG Low Frequency, Standardized Spindle and Theta Amplitude and Ripples. ***A***: EEG amplitudes modulation in frequencies 1-40Hz as a function of time. ***B***: The amplitude modulation of the spindle band (14-18Hz) across time for physical exercise (red line) and rest (blue line). Shaded areas represent the standard error across subjects. The gray shading indicates the interval displaying significant differences between rest and PE.

#### III Ripple activity

Ripples were first observed in rodent hippocampus^9^ and later in humans in a frequency range between 80-150 Hz^10–12^. We detected ripple events in the ongoing MEG signal in accord with previous studies ^4,13,14^ (see ***Fig. 3A*** and ***Methods***). Ripples in MEG resembled the time-frequency representation (mean duration = 31.91 msec, SD = 10.82 msec; ripple frequency = .125 Hz, SD = .098 Hz) observed in intracranial studies ^4,13,14^ (see ***Fig. 3B***).

**Fig. 3.**
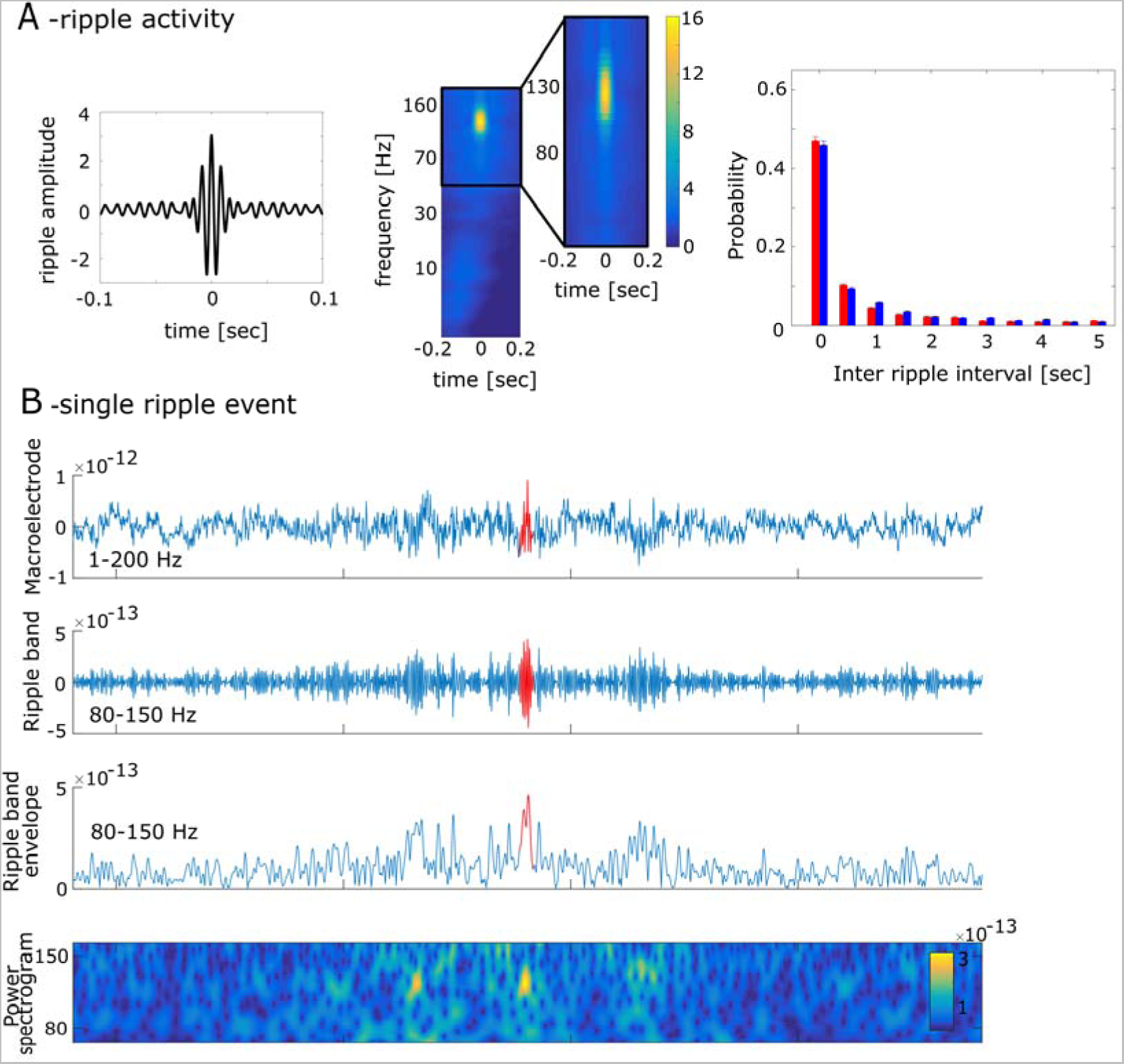
high frequency ripple activation in MEG. ***A*** (left): The grand-averaged signal of ripple activity. (middle): Time-frequency plot of ripple activity. (left): The probability of the interval time between two ripples across subjects. The error bars represent the standard deviation. ***B***: An example of ripples detected in our analysis. The upper line shows the broad band signal between 1 and 200 Hz. The second time series shows the same signal in the ripple band. The third time series shows the Hilbert transform of the ripple band activity. The last graphic shows the time frequency representation of the signal filtered between 80 and 150Hz.

##### Topographical likelihood distribution of ripples

We defined the topographical distribution of ripple events across the entire experiment. For each sensor, we determined the ripple likelihood as the average number of ripples across all trials, including targets and standards, leading to a likelihood value for each of the 102 MEG sensors and each participant. We then correlated the spatial likelihood distribution between all possible pairs of participants. This resulted in 210 correlation values both for rest and PE. We tested whether the distribution of ripples differed between PE and rest (see ***Fig. 4A***). Following PE, pairs of participants exhibited stronger topographical consistency in ripple distribution (mean r_rest_ = 0.37; mean r_PE_ = 0.56; t_209_ = 8.07; p < 0.0001) with sensors covering the MTL region showing the highest ripple likelihood.

**Fig. 4.**
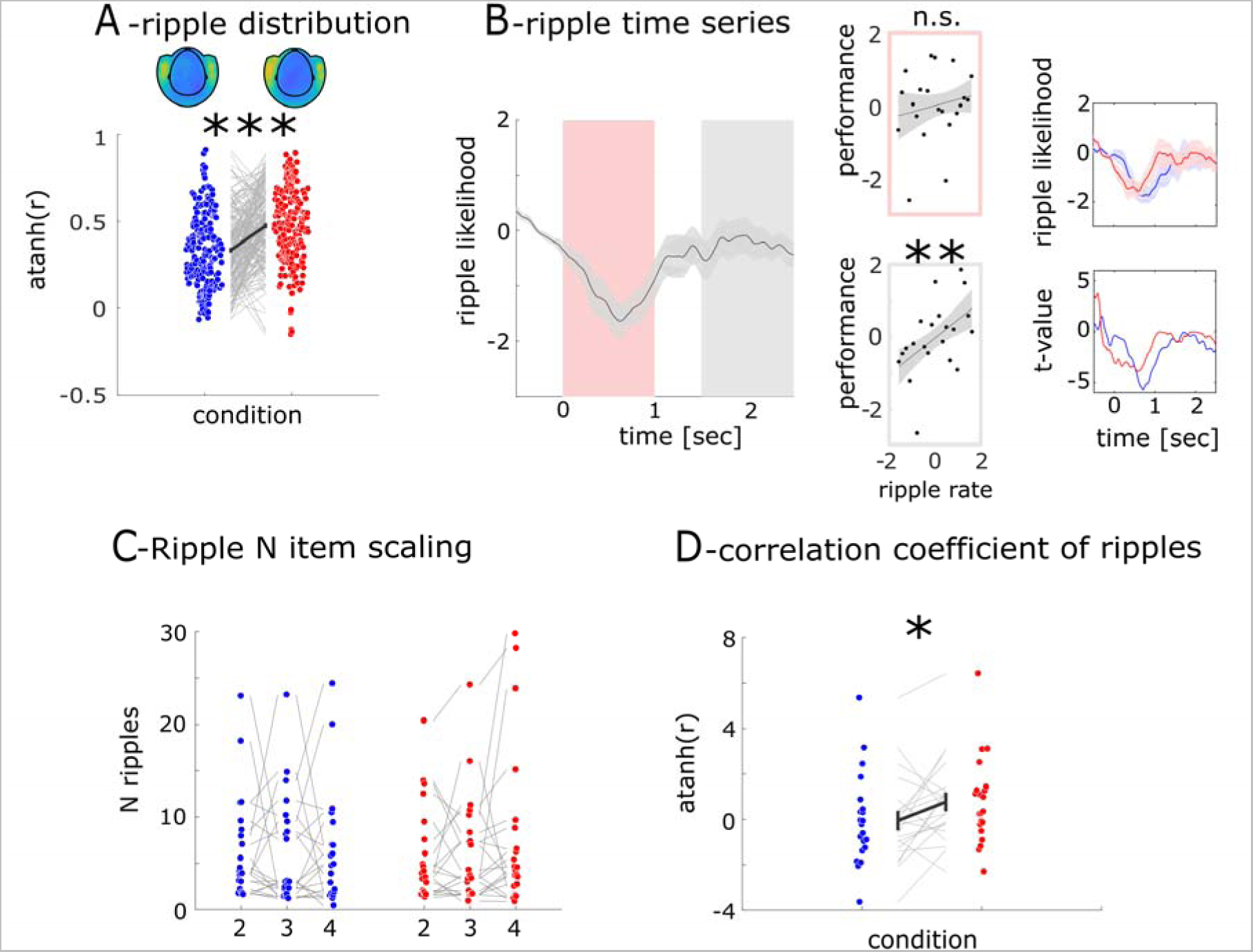
Ripple modulation. ***A*** (upper): Topography map of ripple likelihood across MEG sensors for rest (left) and PE (right). (lower): Correlation coefficients of ripple distributions between pairs of subjects. Higher correlations coefficients following PE indicate a more consistent spatial structure of ripples likelihood across subjects ***B*** (left): Time series of ripple likelihood. (middle): linear regression between individual memory capacity and ripple likelihood in segmented intervals. (right): The ripple likelihood time series from grand-averaged and different sessions, along with the corresponding t-values. ***C***: Ripple count across N-back conditions displayed individually. ***D***: Correlation coefficients for rest (blue) and PE (red) calculated between the ripple likelihood and the 2, 3, 4-back levels.

##### Ripple time series

In the next step, we defined the time course of ripple events around stimulus presentation. We found that the averaged ripple likelihood decreased after stimulus onset (from 280 msec to 970 msec) with a rebound starting around 1.5 sec and peaking around 2 sec (see ***Fig. 4B***).

##### The correlation of ripples with memory capacity

We then tested whether the ripple frequency predicted individual hit rate. We found no correlation in the first second after stimulus presentation (r = 0.17; p = 0.46). However, ripple rate between 1.5-2.5 sec predicted individual target detection rate (r = 0.49; p = .021) (see ***Fig. 4B***). In the first step we compared ripple rate in this interval between conditions but did not find a difference (t_20_ = .43; p = .67). Next, we tested whether the number of ripples is scaled with the number of items to be held in working memory (see ***Fig. 4C***). We conducted a correlation analysis between N-back and ripple rate for each participant leading to a correlation value for each subject. Using directed t-tests under the assumption that r-values are higher than zero (more ripples when more items are held in WM) we found a positive linear scaling of N-back and ripples in the PE session (r_PE_ = 0.77; t_20_ = 1.86; p = 0.0385). No such effect was observed in the rest session (r_Rest_ = −0.05; t_20_ = 0.12; p = 0.452). The N-back ripple correlation in the PE session was stronger than in the rest session (t_20_ = 2.12; p = 0.024, see ***Fig. 4D***).

#### IV MEG-Ripple to EEG-low-Frequency coupling

##### Ripple centered EEG epochs

We tested whether we could find coupling between MEG-ripples and EEG-spindles. Note that we did not find MTL spindle activity in the MEG (see ***Fig. 3A***). We epoched EEG data around MEG-ripple events and found a clear oscillation locked to ripples (see ***Fig. 5A***). This oscillation showed a significant positive amplitude modulation from −14 to 0 msec (t_max_ = 3.24 at −6 msec relative to ripple peak; p = .0041; interval I) and a significant negative amplitude modulation from 10 to 32 msec (t_max_ = 4.08 at 24 msec following ripple peak; p = .0006; interval II) (see ***Fig. 5A***). No such difference could be found for randomized ripple events (interval I: t_max_ = 1.26; p = .57; interval II: t_max_ = .99; p = .32; see ***Fig. 5A***). This pattern can be explained by oscillatory modulation around ripples in the PE session with a stronger modulation in interval II (interval I: t_max_ = 2.9 at −6 msec; p = .011; interval II: t_max_ = 3.7 at 22 msec; p = .0014). No such effect was found in the rest session (interval I: t_max_ = 3.32; p = .003; interval II: t_max_ = 2.2; p = .032)

**Fig. 5.**
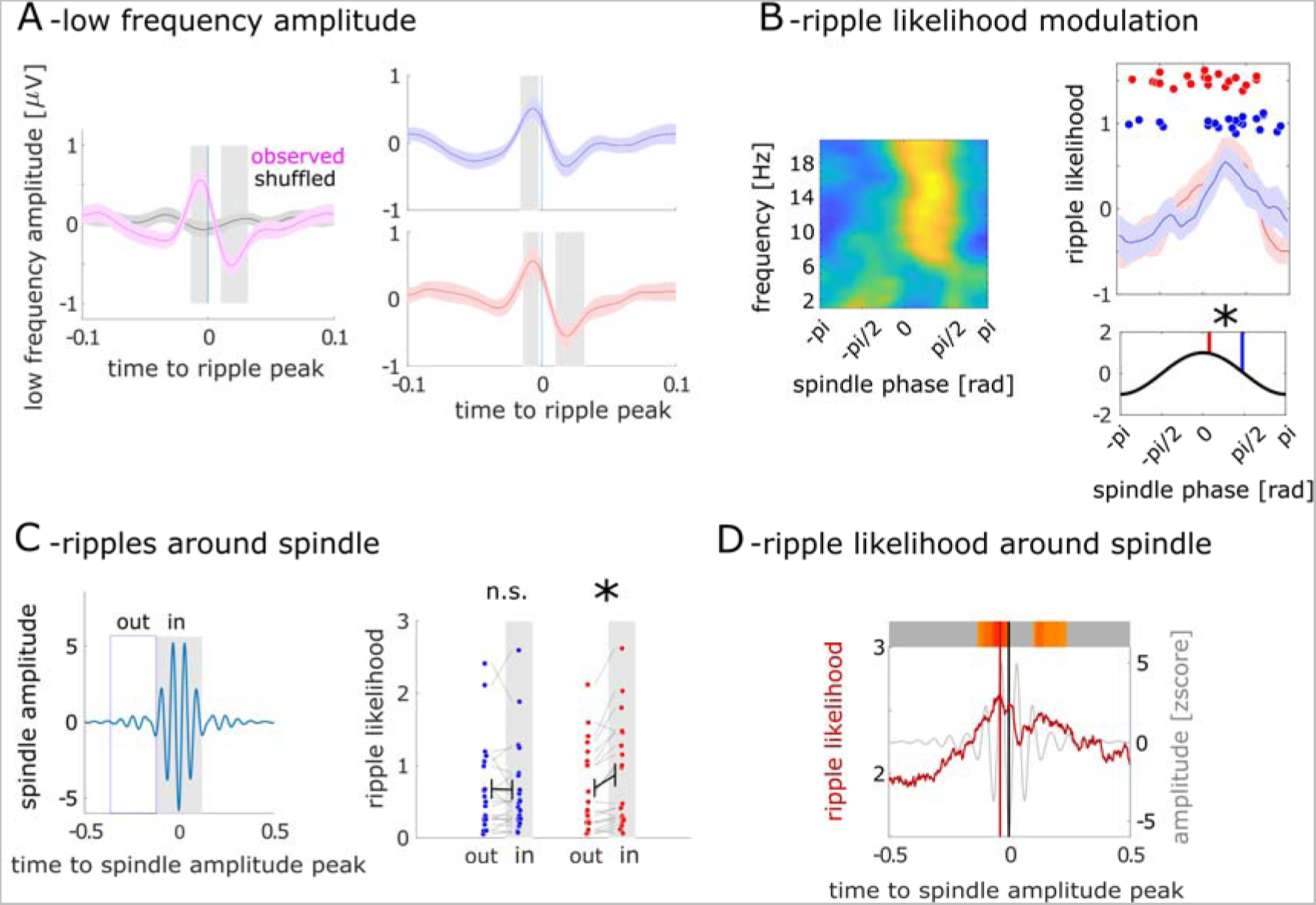
MEG-Ripple and EEG low-frequency coupling results. ***A*** (left): Grand-averaged slow oscillations around the ripple peak compared to the shuffled peak. (right): The slow oscillations around the ripple peak from different sessions. ***B*** (left): Phase distribution of low frequency around the ripple peak. (right upper): The phase distribution from 14-16 Hz from different sessions and the dominant phase from individual subjects. B (right lower): Average dominant phases across sessions. ***C***: Ripple likelihood during spindle (in) and out of spindle (out) phases, plotted individually for PE and rest sessions. Red denotes PE session ripple likelihood, while blue represents ripple likelihood in the rest session. ***D***: Ripple likelihood around the spindle peak. The red time region represents a probability value of p < 0.0001.

##### Ripple-peak to spindle-phase coupling

We then investigated whether ripples are coupled to a specific low-frequency phase. We analyzed the phase of EEG low-frequency activity associated with the ripple peak (see ***Fig. 5B***) and tested whether ripples are differently locked to the spindle band (see ***Fig. 5B***). We found that ripples were locked to different phases φ depending on the session (φ_PE_ = 0.24; φ_rest_ = 1.49;F = 7.05; p = 0.011).

##### Temporal coordination of ripples with spindle events

We examined whether ripples were temporally locked to spindle events by comparing the frequency of ripples outside and within spindle events (see ***Fig. 5C*** and ***Methods*** for a definition of spindle events). In the PE session, we observed more ripples within than outside spindle events (N_in_ = 0.87; N_out_ = 0.69; t_20_ = 3.93; p < .001), an effect not seen in the rest session (N_in_ = 0.67; N_out_ = 0.68; t_20_ =.24; p = .83). In the final step, we investigated when the ripple likelihood reaches its maximum relative to the spindle peak event. We found a peak in the ripple likelihood at approximately 40 msec before the spindle event peak (p < 0.0001)(see ***Fig. 5D***).

## Discussion

We investigated the impact of PE on working memory observing WM performance increases after PE in all N-back conditions. EEG-spindle activity, considered a crucial marker for memory consolidation in sleep research, was heightened in the PE session compared to rest. High-frequency ripples linked to the temporal organization of WM information were detected in MEG sensors covering the medial temporal lobe (MTL) region and the number of ripples predicted individual WM performance. Ripple rate was comparable between PE and rest. However, ripple rate increased with the number of items held in WM. Ripple-spindle coupling also increased following PE.

Moderate PE improved working memory capacity in line with previous studies. Abou Khalil et al.^15^ found a performance increase in a word-recall task during brisk walking compared to sitting. Previous studies found that concurrent moderate exercise led to an improvement in WM accuracy and faster reaction times^16^ and short bouts of exercise improved memory consolidation^17^. In summary, our findings contribute to the growing body of evidence indicating that short-term physical exercise enhances memory processes.

Spindle activity, during non-rapid-eye-movement (NREM) sleep, increases during memory consolidation^4,5^. It has been proposed that spindles play a key role in regulating information flow between the cortex and hippocampus areas, supporting memory consolidation through the reactivation of prior experiences and enhancing structural changes at cortical sites involved in prior learning. The increase in ripples is predictive of individual performance differences ^14,18,19^. The impact of exercise on sleep spindle activity is uncertain, with some studies suggesting that daily repeated exercise can enhance 13-16 Hz spindle power during sleep^20^, while others did not find effects of exercise on spindle activity^21^. We observed spindle activity in both PE and rest sessions, topographically overlapping with sleep spindle findings in previous studies^8^. The authors found spindle amplitude during WM task in alignment with our study demonstrating PE improved working memory performance and increased spindle activity.

Ripple events, considered reactivation patterns, manifest as transient high frequency bursts (~80–150 Hz in humans: mean duration ~ 30 msec). Ripple events were first studied in the hippocampus and more recently identified in extrahippocampal regions^7,22^, particularly during sleep. Numerous studies highlight the crucial role of hippocampal sleep ripples in the initial formation of declarative memory traces, suggesting their involvement in anatomically distributed memory traces^23^. Hippocampo– thalamocortical synchronization during sleep is key to human memory consolidation^24^. A prominent model posits that embedding novel information in the neocortex requires offline reactivation by hippocampal generated ripple events, primarily during slow-wave sleep^18,25^. Recent studies have also reported ripples during waking, influencing emotional memory encoding and discrimination^26^. Ripples in the hippocampus, MTL, and neocortex are reported to mirror neuron firing patterns during cued recall. Awake ripples share similarities in density, frequency, and duration during both waking and sleep^11^. Episodic memory studies in human subjects support the role of the hippocampus and surrounding cortical areas in learning and recalling temporal sequences. We show that MTL ripple rate scales with the number of items held in working memory and predicts individual working memory capacity supporting a reactivation mechanism. Ripples show a specific time course with the ripple rate decreasing during stimulus presentation, rebound around 1 sec, and predict individual performance.

We observed that ripple rate predicted the number of items in working memory post-PE aligning with prior research indicating that ripple rate scales with the complexity of autobiographical memory ^27^. We also identified a correlation between ripple activity and hit performance, consistent with previous results reporting a correlation between ripple rate and successful memory retrieval^18^.

Spindles and high-frequency ripple activity were co-modulated and modulated by PE. Spindle-ripple coupling was initially described during rat slow-wave sleep^28^ and reported in epilepsy patients during sleep using intracranial EEG (iEEG) or parahippocampal foramen ovale (FO) electrodes^7,10,29^. In line with these studies, we found coupling of EEG spindle activity with MEG ripple events in both the PE and rest sessions. However, PE modulated coupling through phase- and temporal alignment with spindle activity, exhibiting a stronger spindle modulation around ripple events. This coupling property is consistent with previous studies showing that ripple rates increase during the ‘waxing’ spindles phase (the raising phase) from the iEEG recording at human MTL regions ^7^. Physical exercise shifted the phase of ripple-coupled spindles, temporally aligning ripples with high-amplitude spindle events. Our results indicate that short-term PE enhances spindle-ripple coupling, contributing to improved memory formation. It is proposed that coupling creates a temporal window for communication between the hippocampus and neocortical regions ^30,31^. The enhanced spindle-ripple coupling and the phase shift observed during PE session may signify enhanced communication among reactivation processes between cortical and subcortical structures enhancing working memory.

## Methods

### Participants

21 participants (13 female, 8 male, mean age = 27.00 years, SD = 5.76 years) participated in the experiment. All participants reported normal or corrected to normal vision and no history of neurological or psychiatric diseases. The recordings took place at the Department of Neurology, Otto-von-Guericke University Magdeburg. We invited participants to two separate MEG sessions with an average 75-day (SD = 65.27) interval between sessions to allow for a within-subject design. They either started with a PE or rest session counterbalanced across participants.

### Procedure

The stimulus presentation and experimental control were carried out using Matlab R2009a (Mathworks, Natick, USA) and the Psychophysics Toolbox ^32^. The screen color was set to gray, while the stimuli and instructions were displayed in black. The experiment consisted of 12 blocks, each comprising 100 trials of an N-back task. Target items were defined as items that matched the item presented in either N=2, N=3, or N=4 trials before the current trial requiring subjects to hold a varying number of items in their working memory. Each block consisted of 30 targets and 70 standards. Stimuli included 26 letters of the Latin alphabet, with targets and standards randomly selected. The experiment started with a practice session involving N=2 and N=4 back tasks, which only included 6 targets and 14 standards, under identical presentation settings as the main experiment. Each block began with a 2-minute phase before the actual visual stimuli were presented. Participants either used the pedal trainer or remained seated, depending on the session (rest vs. PE). Each block lasted 252 seconds. Each trial started with a fixation cross presented for 500 msec, centered on the screen. The participants were instructed to focus the fixation cross during the entire experiment to minimize eye movements. Subsequently, the fixation cross was replaced by a letter stimulus for 500 ms, followed by the reappearance of the fixation cross for 2000±100 ms.

Participants were instructed to respond as quickly and accurately as possible. They were instructed to use their index finger to respond to standards and their middle finger to respond to targets. The responses were provided with the left and right hands, which were alternated every three blocks. The pedal trainer was purposefully designed in-house for the MEG recording. All the materials were thoroughly tested ensuring the absence of any magnetic components. The device allows for the adjustment of both angle and height, customizing it to fit an individual participant’s proportions. Its design permits forward and backward movements of both feet individually. The participants were instructed to pedal at a moderate speed, adjusting their pace individually, akin to a walking motion.

### MEG Recording

Participants were equipped with metal-free clothing and seated in a dimmed, magnetically shielded recording booth. Stimuli were presented via rear projection onto a semi-transparent screen with an LCD projector (DLA-G150CLE, JVC, Yokohama, Japan) that was positioned outside the booth. The screen was placed at a viewing distance of 100 cm in front of the participants. Responses were given with the left and right hands via an MEG compatible LUMItouch response system (Photon Control Inc., Burnaby, DC, Canada). Acquisition of MEG data was performed in a sitting position using a whole-head Elekta Neuromag TRIUX MEG system (Elekta Oy, Helsinki, Finland), containing 102 magnetometers and 204 planar gradiometers. The sampling rate was set to 2000 Hz. EEG was recorded with 30 passive electrodes. The right mastoid was used as a reference electrode. Vertical EOG was recorded using one surface electrode above and one below the right eye. For horizontal EOG, one electrode on the left and right outer canthus was used. Preparation and recordings took about 3 hours.

### Preprocessing and Artifact Rejection

Maxwell filtering was applied to reduce external noise and both MEG and EEG data were down sampled to 500 Hz. We used Matlab 2016b (Mathworks, Natick, USA) for analysis. We included the 102 magnetometers in our analyses but no gradiometers. All filtering (see below) was done using zero phaseshift IIR filters (fourth order butterworth filter; filtfilt.m in Matlab). First, we filtered the data between 1 and 200 Hz. Then, we notchfiltered the data to discard line noise (50Hz and its 2nd and 3rd harmonic). To discard trials of excessive, nonphysiological amplitude, we used an individual threshold for each subject. Both for MEG and EEG we calculated the mean variance across time and channels for each trial. Trials with a variance exceeding 4 standard deviations were excluded. We then visually inspected all data and excluded epochs exhibiting excessive muscle activity, as well as time intervals containing artifactual signal distortions, such as signal steps or pulses.

EEG data were re-referenced to the left mastoid and band-pass filtered between 1 to 40 Hz. To eliminate eye movement artifacts, we applied Independent Component Analysis (ICA), which was computed with the FastICA package for MATLAB (Version 2.5). Resulting MEG and EEG time series were used to characterize ripple band dynamics (see below) and low frequency activity, respectively, over the time course of visual target detection. Both MEG and EEG data were epoched into trials ranging from −2 to 3 sec (sufficiently long to prevent edge effects).

### Statistical Analysis

In this study, we aimed to explore the effects of physical exercise on working memory and the associated changes in brain states, specifically focusing on spindle and ripple activity as well as their coupling effect. First, we compared target detection performance and reaction times between PE and rest across various N-back conditions (I - Behavioral Results). In the next step, we examined EEG spindle activity (II - Spindle activity). Subsequently, we employed MEG signals to compute ripple activity and investigated correlations between task performance, memory load with ripple activity (III - Ripple activity). Lastly, we tested for spindle-ripple coupling and differences between rest and PE (IV - Spindle-Ripple Coupling).

#### I Behavioral performance

We evaluated target detection accuracy based on hit rate (DA; percentage of target response when a target was present), false alarms (FA; percentage of target response when no target was presented), and reaction times (RT). We compared the impact of PE and rest on behavioral measures using a two-way ANOVA with the factors ***motion*** (physical exercise vs. rest) and ***N-back*** (2-back, 3-back, 4-back) separately for DA, FA and RT.

#### II EEG spindle activity

It is assumed that spindle activity establishes a timeframe in which ripples occur^7^. In the EEG, spindle activity oscillations generated within thalamocortical loops can predominantly be measured at fronto-central electrodes^8^. To verify this pattern, we tested which low-frequency bands exhibit strongest modulation following stimulus onset in a set of four fronto-central electrodes (Fz, Cz, Fc1, and Fc2). These channels were also utilized for subsequent spindle and coupling analyses. To calculate the time-frequency representation, we band-pass filtered the MEG signal at 21 exponentially increasing center frequencies (ranging from 1 to 40 Hz), each with a bandwidth of 10% around the center frequency bands. The resulting time series were Hilbert-transformed, epoched around stimulus onsets (from −1 to 2 sec) (see ***Fig. 2A***), baseline-corrected by subtracting the average activity in the 200 msec prior to stimulus onset from each time point, and averaged across all trials. The resulting time series were z-scored with respect to a baseline period (−200 msec prior to stimulus onset) for a better comparison across frequencies. For this we firstly calculated the mean and standard deviation of time series in the baseline period and then subtracted the mean from the time series and divided them by the standard deviation at each time point. The following z-score analysis with baseline correction was computed with the same procedure. The time-frequency representation confirmed clear spindle activity. We then extracted EEG-spindle activity in the frequency band showing highest amplitudes. We band-pass filtered the EEG signal between 14 and 18 Hz. Subsequently, we determined the analytic amplitude Af(t) by Hilbert-transforming the signal maintaining the same baseline correction settings. The resulting spindle activity was averaged across all trials separately for rest and PE, then we z-scored the spindle activity with a baseline correction period from −200 msec to stimulus onset (see ***Fig. 2B***). To investigate the disparity in spindle activity between the rest and PE under different time courses, we conducted a t-test to analyze the difference between two sets of data, taken at intervals of 0.1 seconds after the stimulus from 0 to 2.5 seconds.

#### III Ripple activity

Ripple events were detected in the ongoing MEG signal in the following way ^13,14,33^. Ripples were first observed in rodent hippocampus which occurred around 100-200 Hz ^9^, while ripples in humans, which share many characteristics with it, occur at frequencies around 80-150 Hz^10,11^. Hence, the MEG signal was band-pass filtered between 80 and 150 Hz. We obtained the analytic amplitude A(t) by applying a Hilbert transform to the filtered time series. To avoid that potential filtering artifacts affect ripple detection, A(t) was reset to zero during the initial and final 100 msec of each trial. To establish a distribution of ripple events throughout the entire experiment, we concatenated all trials and computed the z-score Z(t) separately for each MEG sensor. Ripple events were identified as peaks in the Hilbert signal that exceed 2.5 SD ^33^ above mean, extend over 20 ms up to 500 msec (equivalent to 3 cycles at 150 Hz and 40 cycles at 80 Hz), and were separated by at least 20 ms. Events exceeding 9 SD above mean were excluded. We then retained only ripple events with at least three peaks that exceeded 2.5 SD above the mean, with at least one peak designated as dominant (prominence ≥ 20% above neighboring peaks).

##### Time frequency representation of ripple events

We tested whether ripples in MEG resemble the time-frequency representation in intracranial studies ^13,14,33^ in two steps. First, we calculated the grand average ripple shape. To this end, we epoched the raw MEG signal (filtered between 1 and 200 Hz) around the ripple peak events. Within subjects we averaged all resulting ripple-centered time series leading to a ripple shape for each subject (***Fig. 3A***). Second, we calculate the time frequency representation by band-pass filtering the MEG signal at 38 exponentially increasing center frequencies (between 1 and 200 Hz) each with a band width of 10% around the center frequency bands. We obtained the analytic amplitude ***A(t)*** by Hilbert-transforming the filtered time series. Then we epoched the frequency specific ***A(t)*** around ripple events (***Fig. 3A***) and averaged across epochs. The resulting time series were z-scored as to a baseline period (−1.0 and −0.2 sec) before the ripple event.

##### Frequency distribution of ripples

First, we determined the topographical distribution of ripple events across the entire experiment. For each sensor, we calculated the ripple likelihood as the average number of ripples across all trials leading to ripple rate for each of the 102 MEG sensors and each participant. We tested whether the distribution of ripples differed between PE and rest (***Fig. 4A***). Specifically, we tested whether the topographical ripple distribution becomes more stable across subjects with PE. To do this, we utilized the individual vectors of the ripple likelihood distribution across the 102 MEG sensors for each participant, both during PE and rest. We then correlated the likelihood distribution between all possible pairs of participants. This resulted in 210 correlation values both for rest and PE. The resulting correlation coefficients were converted into a metric distribution by calculating its inverse hyperbolic tangent and compared using a t-test. In the following steps, we limited the analysis to the MEG sensors with the highest number of ripples, which were four sensors positioned bilaterally covering the MTL region. The ripple peak time points were used to calculate the grand average ripple shape and the time frequency representation, and coupling with EEG spindle activity. For all the calculations, we chose a two-second window centered around the ripple events (from −1.0 to + 1.0 sec).

##### Ripple time series

In the next step, we defined the time course of ripple events around stimulus presentation (***Fig. 4B***). We summed the number of ripples detected across the four most informative sensors and all trials in each participant. This resulted in a discrete time series of zeros and ones (indicating a ripple event) for each participant. We then convolved the resulting time series in the following manner. Within a moving time window of 200 msec, we summed ripple events. This yielded a continuous time series R for each participant. Then, we standardized R (z-score) according to baseline activity.

##### The correlation of ripples with memory capacity

Next, we tested whether the ripple frequency predicts individual hit rate. The ripple likelihood time series showed a continuous decrease over one second after stimulus onset with a following rebound. Therefore, we tested whether the initial decrease or the subsequent rebound is predictive of differences in performance. For each participant, we determined the mean ripple likelihood from 0 to 1 sec and from 1.5 to 2.5 sec after stimulus onset (see ***Fig. 4D***). To investigate whether ripples correlated with individual performance, we averaged target detection performance across all blocks and physical activity sessions (rest and PE). We then determined the rank of performance across all participants and correlated this with the rank of individual ripple likelihood using Pearson correlation. In the next step, we investigated whether the number of ripples provided information about the number of items that need to be retained in working memory. We correlated the ripple frequency in different N-back conditions with N-back (2-3-4, see ***Fig. 4C,D***). For each participant, we determined the mean ripple frequency separately for each N-back condition. We then correlated these three resulting values with the 3 N-values separately for the rest and PE session. As a result, we obtained Pearson’s r value for each participant, both for the rest and PE session. These values were translated into a metric measure by calculating their inverse hyperbolic tangent and were compared between rest and PE using a t-test across participants.

#### IV MEG-Ripple to EEG-low-Frequency coupling

##### Ripple centered EEG epochs

Ripple and spindle activity have been observed in the human medial temporal lobe and ripple-spindle coupling is described as signature of memory consolidation in rodents^34,35^. Analogous findings in humans can be found during sleep^36^. Here we tested whether we could find a similar coupling pattern during wakefulness with non-invasive measurements. In the first step, we tested whether a modulation of EEG activity occurs simultaneously with ripple events (***Fig. 5A right***). We epoched the filtered EEG signal (1-25 Hz) at the fronto-central channels around the ripple events. We identified ripples as outlined above and segmented the EEG data into epochs of length 2 sec (−1 to 1 sec) centered to the peak of the ripple. Resulting time series for each ripple event and each channel were averaged within subjects. This yielded a time series RS_(t)_ for each participant. We then standardized RS_(t)_ (z-score) according to baseline activity. First, we calculated the mean and standard deviation of the RS_(t)_ in the baseline period. Here, we chose the time interval between −1.0 and −0.5 sec before the ripple event as the baseline since we expect spindle events to be longer in duration than ripple events. Finally, we subtracted the mean from the RS_(t)_ at each time point and divided the RS_(t)_ at each time point by the standard deviation. We compared the spindle amplitude of the ripple centered EEG at each time point against 0 using a t-test. In the next step, we constructed surrogate EEG signals which were also tested against 0. The surrogate signals were constructed by centering the EEG signals to random time points matching the number of ripples in each subject. In this analysis, we shifted the ripple time series for each participant over time. This allows for maintaining both the number of ripple events and their intervals consistently. With these new surrogate ripple events, we repeated the analysis described above (***Fig. 5A left***).

##### Ripple Spindle coupling

In the next step, we investigated whether ripples are coupled to a specific low-frequency phase (see ***Fig 5B***). Phase-amplitude cross-frequency coupling (PAC) is a mechanism that has been proposed to coordinate the timing of neuronal firing within local neural networks^37^. We utilized conventional cross-frequency coupling metrics^38,39^ to test for coupling of ripple events to low frequency bands in EEG channels. Generally, we determined at which phase of the low-frequency activity ripples most frequently occur. To do this, we initially determined the phase angle at each time point for 20 frequency bands ranging from 1 to 20 Hz each. First, we band-pass filtered the EEG signal at these center frequencies each with a width of 2 Hz. We then extracted the instantaneous phase information using the Hilbert transformation. We calculated the instantaneous phase of the low frequency activity for each EEG channel time series. We divided each low frequency cycle separately in 50 equally spaced bins ranging from –π to π and summed the number of ripples within a 45-degree window centered on every phase bin ^40^. The resulting ripple event histograms – each containing 50 values – were averaged, separately for each low frequency band (see ***Fig 5B right***). We then averaged the dominant phase accross different subjects with (circ_mean in Circular Statistics Toolbox) and compared the phases with parametric Watson-Williams multi-sample test (circ_wwtest in Circular Statistics Toolbox; see ***Fig 5B right lower***).

##### Temporal coordination of ripples with spindle events

In the previous steps, we examined whether ripple peaks are generally coupled with spindle activity. However, spindle activity is not a uniform oscillation but is characterized by specific high-amplitude bouts^31^. In the next step, we examined whether ripples are temporally locked to spindle events by comparing the frequency of ripples outside and within spindle events. We defined spindle events as follows: first, we filtered the EEG signal in the spindle frequency (13-20 Hz) and obtained the analytic amplitude ***A***_(t)_. The resulting time series was z-scored, and peaks with z>4 were detected. Note, that the peak of ***A***_(t)_ is not necessarily the peak/trough of the spindle oscillation. Given that ripples are aligned with a specific phase (see ***Results***) of the spindle activity we centered spindle events to the troughs of spindle oscillation. Hence, we epoched the data (−2 to 2 sec) around each detected peak in ***A***_(t)_. We then detected the trough of the spindle oscillation closest to the peak of ***A***_(t)_. This timepoint was used as the center of the spindle event. Subsequently, we epoched the ripple time series around these spindle events. These ripple epochs RE_t_ again contain zeros and ones (indicating a ripple peak event). We then tested when ripples occurred relative to individual spindle events. For this, we defined two temporal intervals. The within spindle interval was defined as the time range from −0.1 to 0.1 sec (200 msec ~ 3 cycles), while the outside spindle interval was defined as the time range from −0.3 to −0.1 sec. Separately for both intervals, we summed the ripples over all spindle events for each subject (see ***Fig 5C***). This results in a ripple event rate both for the within and outside interval for each subject. This was done separately for the PE and rest session. The resulting ripple event rates were then compared using a t-test in each session.

Finally, we tested when ripple likelihood peaks relative to spindle peak events (see ***Fig 5D***). A peak of ripple likelihood before the respective spindle peak event might indicate that ripples trigger spindles. In contrast, a peak of ripple likelihood following the spindle peak event might indicate that spindle events drive ripples. First, we summed ripple epochs (RE_t_) across all spindle events for each participant, creating a time series of ripple likelihood around the spindle event peak for each participant. These individual time series were then averaged across subjects. This averaged ripple rate was compared against a surrogate distribution. To construct this distribution, we shifted the summed RE_t_ around spindle event peaks for each participant over time and averaged the resulting time series across participants. This process was repeated 1000 times, resulting in 1000 surrogate ripple event series. We determined the likelihood of ripple events at each time point relative to the surrogate distribution at each time point by estimating the probability density functions (pdf.m in Matlab). This yielded a probability value ***p*** for each time point around the spindle event peak.

### Data availability

All data supporting the results presented in the results section and figures can be accessed on github (https://github.com/SDuerschmid/PhysicalExerciseWorkingMemory) upon publication.

## Author contributions

S.D. conceived and designed the experiment. X.C. collected the MEG data. X.C., B.A., P.S., C.R., A.S., T.W., and S.D. analyzed the data, X.C., B.A., P.S., C.R., R.T.K., A.S., T. W., and S.D. interpreted the data. X.C., R.T. K., and S.D. wrote the manuscript.

## Competing Interest Statement

no competing interests

## Notes

https://github.com/SDuerschmid/PhysicalExerciseWorkingMemory

### Competing Interest Statement

The authors have declared no competing interest.

## References

1. Roig, M., Nordbrandt, S., Geertsen, S. S. & Nielsen, J. B. The effects of cardiovascular exercise on human memory: A review with meta-analysis. Neuroscience and Biobehavioral Reviews vol. 37 1645–1666 Preprint at 10.1016/j.neubiorev.2013.06.012 (2013).

2. van Praag, H., Christie, B. R., Sejnowski, T. J. & Gage, F. H. Running enhances neurogenesis, learning, and long-term potentiation in mice Enhanced Reader. PNAS 96, 13247–13431 (1999).

3. Voss, M. W. et al. Exercise and Hippocampal Memory Systems. Trends in Cognitive Sciences vol. 23 318–333 Preprint at 10.1016/j.tics.2019.01.006 (2019).

4. Latchoumane, C. F. V., Ngo, H. V. V., Born, J. & Shin, H. S. Thalamic Spindles Promote Memory Formation during Sleep through Triple Phase-Locking of Cortical, Thalamic, and Hippocampal Rhythms. Neuron 95, (2017).

5. Schreiner, T., Petzka, M., Staudigl, T. & Staresina, B. P. Endogenous memory reactivation during sleep in humans is clocked by slow oscillation-spindle complexes. Nat Commun 12, (2021).

6. De Vries, W. R., Bernards, N. T. M., De Rooij, M. H. & Koppeschaar, H. P. F. Dynamic exercise discloses different time-related responses in stress hormones. Psychosom Med 62, (2000).

7. Staresina, B. P., Niediek, J., Borger, V., Surges, R. & Mormann, F. How coupled slow oscillations, spindles and ripples coordinate neuronal processing and communication during human sleep. Nature Neuroscience 2023 1–9 (2023) doi:10.1038/s41593-023-01381-w.

8. Petzka, M., Chatburn, A., Charest, I., Balanos, G. M. & Staresina, B. P. Sleep spindles track cortical learning patterns for memory consolidation. Current Biology 32, (2022).

9. Buzsáki, G., Horváth, Z., Urioste, R., Hetke, J. & Wise, K. High-frequency network oscillation in the hippocampus. Science (1979) 256, (1992).

10. Clemens, Z. et al. Temporal coupling of parahippocampal ripples, sleep spindles and slow oscillations in humans. Brain 130, (2007).

11. Dickey, C. W. et al. Cortical ripples during NREM sleep and waking in humans Title: Cortical ripples during NREM sleep and waking in humans 1 2 Abbreviated title: Cortical ripples in humans 3 4. Cite as: J. Neurosci 10, 4 (2022).

12. Tong, A. P. S., Vaz, A. P., Wittig, J. H., Inati, S. K. & Zaghloul, K. A. Ripples reflect a spectrum of synchronous spiking activity in human anterior temporal lobe. Elife 10, (2021).

13. Staresina, B. P. et al. Hierarchical nesting of slow oscillations, spindles and ripples in the human hippocampus during sleep. Nat Neurosci 18, 1679–1686 (2015).

14. Norman, Y. et al. Hippocampal sharp-wave ripples linked to visual episodic recollection in humans. Science (1979) 365, eaax1030 (2019).

15. Abou Khalil, G., Doré-Mazars, K., Senot, P., Wang, D. P. & Legrand, A. Is it better to sit down, stand up or walk when performing memory and arithmetic activities? Exp Brain Res 238, (2020).

16. Quelhas Martins, A., Kavussanu, M., Willoughby, A. & Ring, C. Moderate intensity exercise facilitates working memory. Psychol Sport Exerc 14, (2013).

17. Most, S. B., Kennedy, B. L. & Petras, E. A. Evidence for improved memory from 5 minutes of immediate, post-encoding exercise among women. Cogn Res Princ Implic 2, (2017).

18. Vaz, A. P., Inati, S. K., Brunel, N. & Zaghloul, K. A. Coupled ripple oscillations between the medial temporal lobe and neocortex retrieve human memory. Science (1979) 363, (2019).

19. Norman, Y., Raccah, O., Liu, S., Parvizi, J. & Malach, R. Hippocampal ripples and their coordinated dialogue with the default mode network during recent and remote recollection. Neuron 109, 2767–2780.e5 (2021).

20. Aritake-Okada, S. et al. Diurnal repeated exercise promotes slow-wave activity and fast-sigma power during sleep with increase in body temperature: A human crossover trial. J Appl Physiol 127, (2019).

21. Frisch, N. et al. An acute bout of high-intensity exercise affects nocturnal sleep and sleep-dependent memory consolidation. J Sleep Res (2023) doi:10.1111/jsr.14126.

22. Skelin, I. et al. Coupling between slow waves and sharp-wave ripples engages distributed neural activity during sleep in humans. Proc Natl Acad Sci U S A 118, (2021).

23. Buzsáki, G. Hippocampal sharp wave-ripple: A cognitive biomarker for episodic memory and planning. Hippocampus 25, 1073–1188 (2015).

24. Geva-Sagiv, M. et al. Augmenting hippocampal–prefrontal neuronal synchrony during sleep enhances memory consolidation in humans. Nature Neuroscience 2023 26:6 26, 1100–1110 (2023).

25. Liu, A. A. et al. A consensus statement on detection of hippocampal sharp wave ripples and differentiation from other fast oscillations. Nature Communications vol. 13 Preprint at 10.1038/s41467-022-33536-x (2022).

26. Zhang, H. et al. Awake ripples enhance emotional memory encoding in the human brain Enhanced Reader. Nat Commun 15, (2024).

27. Chen, Y. Y. et al. Stability of ripple events during task engagement in human hippocampus. Cell Rep 35, (2021).

28. Siapas, A. G. & Wilson, M. A. Coordinated interactions between hippocampal ripples and cortical spindles during slow-wave sleep. Neuron 21, (1998).

29. Clemens, Z. et al. Fine-tuned coupling between human parahippocampal ripples and sleep spindles. European Journal of Neuroscience 33, (2011).

30. McKenzie, S., Nitzan, N. & English, D. F. Mechanisms of neural organization and rhythmogenesis during hippocampal and cortical ripples. Philosophical Transactions of the Royal Society B: Biological Sciences vol. 375 Preprint at 10.1098/rstb.2019.0237 (2020).

31. Helfrich, R. F. et al. Bidirectional prefrontal-hippocampal dynamics organize information transfer during sleep in humans. Nature Communications 2019 10:1 10, 1–16 (2019).

32. Brainard, D. H. The psychophysics toolbox. Spat Vis 10, 433–436 (1997).

33. Ngo, H. V., Fell, J. & Staresina, B. Sleep spindles mediate hippocampal-neocortical coupling during long-duration ripples. Elife 9, (2020).

34. Xia, F. et al. Parvalbumin-positive interneurons mediate neocortical-hippocampal interactions that are necessary for memory consolidation. Elife 6, (2017).

35. Diekelmann, S. & Born, J. The memory function of sleep. Nat Rev Neurosci 11, 114– 126 (2010).

36. Ramadan, W., Eschenko, O. & Sara, S. J. Hippocampal sharp wave/ripples during sleep for consolidation of associative memory. PLoS One 4, (2009).

37. Canolty, R. T. & Knight, R. T. The functional role of cross-frequency coupling. Trends Cogn Sci 14, 506–515 (2010).

38. Canolty, R. T. et al. High Gamma Power Is Phase-Locked to Theta Oscillations in. 313, 1626–1629 (2006).

39. Szczepanski, S. M. & Knight, R. T. Insights into Human Behavior from Lesions to the Prefrontal Cortex. Neuron 83, 1002–1018 (2014).

40. Helfrich, R. F. et al. Neural Mechanisms of Sustained Attention Are Rhythmic. Neuron 99, 854–865.e5 (2018).

